# The intracellular amastigote of *Trypanosoma cruzi* maintains an actively beating flagellum

**DOI:** 10.1101/2022.11.23.517661

**Authors:** Madalyn M. Won, Timothy Krüger, Markus Engstler, Barbara A. Burleigh

## Abstract

Throughout its complex life cycle, the uniflagellate parasitic protist, *Trypanosoma cruzi*, adapts to different host environments by transitioning between elongated motile extracellular forms and non-motile intracellular amastigote forms that replicate in the cytoplasm of mammalian host cells. Despite their name, intracellular *T. cruzi* amastigotes retain a short flagellum that extends beyond the opening of the flagellar pocket with access to the extracellular milieu. Contrary to the long-held view that the *T. cruzi* amastigote flagellum is inert, we now report that this organelle is motile and displays quasiperiodic beating inside mammalian host cells. Kymograph analysis determined an average flagellar beat frequency of ~0.7 Hz for intracellular amastigotes. Similar beat frequencies were measured in extracellular amastigotes following their isolation from host cells. Inhibitor studies reveal roles for parasite mitochondrial respiration and intracellular calcium availability in modulating flagellar beat in *T. cruzi* amastigotes. Together, these findings demonstrate that flagellar motility is an intrinsic property of *T. cruzi* amastigotes and suggest that this organelle may play an active role in the parasite infection process. To our knowledge, this is the first record of an intracellular eukaryotic flagellum beating within another eukaryotic cell.

## Introduction

*Trypanosoma cruzi* is the causative agent of human Chagas disease, which is associated with significant morbidity and mortality in Latin America (Bern, 2015; Stanaway & Roth, 2015). The ability of *T. cruzi* to establish intracellular residence in mammalian host cells and to persist in diverse tissues in the host are critical factors underlying disease progression. In mammals, motile trypomastigote stages of the parasite actively penetrate host cells to establish intracellular infection (Fernandes & Andrews, 2012). Once inside, trypomastigotes transform into a non-motile, ‘amastigote’ stage that replicates freely in the host cell cytoplasm (Andrews et al., 1990). Accompanying the loss of motility in the *T. cruzi* intracellular amastigote is a dramatic shortening of the parasite flagellum (Andrews et al., 1987). Often described as a remnant and assumed to be inert (Gardener, 1974; Kelly et al., 2020; Kohl & Bastin, 2005; Kurup & Tarleton, 2014), the *T. cruzi* amastigote flagellum has garnered little attention. However, our previous report of contact occurring between the flagellum of intracellular *T. cruzi* amastigotes and host mitochondria has prompted speculation that the amastigote flagellum may play an active role in infection (Lentini et al., 2018). This observation brings to mind the way the related trypanosomatid, *Leishmania mexicana*, forms a tight interaction between the vacuole-bound amastigote flagellar tip and the parasitophorous vacuole membrane (Gluenz et al., 2010). Thus, yet undiscovered functions of an amastigote flagellum may be conserved across trypanosomatid species.

In addition to propelling motile life stages, trypanosomatid flagella have recognized sensory capabilities that enable adaptation to changing environments (Kelly et al., 2020; Lopez et al., 2015; Oberholzer et al., 2011; Rodriguez-Contreras et al., 2015). In the extracellular African species, *Trypanosoma brucei*, flagellar adenylate cyclases and cAMP-dependent signaling pathway proteins are discussed as critical regulators of transmission and immune evasion (Bachmaier et al., 2022; Lopez et al., 2015; Oberholzer et al., 2015; Salmon, 2018). Furthermore, the flagellar-localized glucose transporter LmxGT1 is thought to act as a glucose sensor for *Leishmania mexicana* promastigotes and amastigotes (Rodriguez-Contreras et al., 2015). Because the *T. cruzi* amastigote flagellum has been found to interact with the host mitochondria, we hypothesized that it might likewise have a comparable sensory role in the mammalian host cell. On this basis, we endeavored to examine the dynamics of the interaction between the *T. cruzi* amastigote flagellum and the host mitochondria and uncovered unexpected flagellar motility instead.

## Results and Discussion

### *T. cruzi* amastigotes have motile flagella

To facilitate live imaging of the *T. cruzi* amastigote flagellum, we generated a transgenic parasite line that stably expresses flagellar-localized SMP1-1-GFP (**Fig 1A**). Similar to the well-characterized SMP1 ortholog in *Leishmania*, which is targeted to the inner leaflet of the flagellar membrane (Tull et al., 2010), SMP1-1-GFP localized primarily to the flagellum in *T. cruzi* amastigotes (**Fig 1A**). Previous fluorescence confocal and super-resolution imaging of fixed *T. cruzi-*infected fibroblasts revealed that the flagella of a high proportion of intracellular amastigotes were in close proximity/contact with host mitochondria (Lentini et al., 2018). To examine the dynamics of this interaction, we performed live fluorescence confocal imaging of SMP1-1-GFP *T. cruzi* amastigotes in fibroblasts expressing mitochondrial COX8-mCherry (**Fig 1B; Supplemental video 1**). As illustrated in time-lapse images (**Fig 1B**), the amastigote flagellum appears to be highly dynamic; its position changes visibly within the ~1-sec interval between each frame. Time-lapse videos of the untagged parental (Tulahuen) strain of *T. cruzi* (**Supplemental Video 2**) confirm that amastigote flagellar movement is not due to ectopic expression of SMP1-1-GFP. Amastigote flagellar movement was also observed in genetically diverse *T. cruzi* strains (**Supplemental Videos 3-5**) and in fibroblasts derived from a different host species (mouse, **Supplemental Video 6**). These observations support the conclusion that flagellar movement is a universal property of intracellular *T. cruzi* amastigotes.

**Figure 1:**
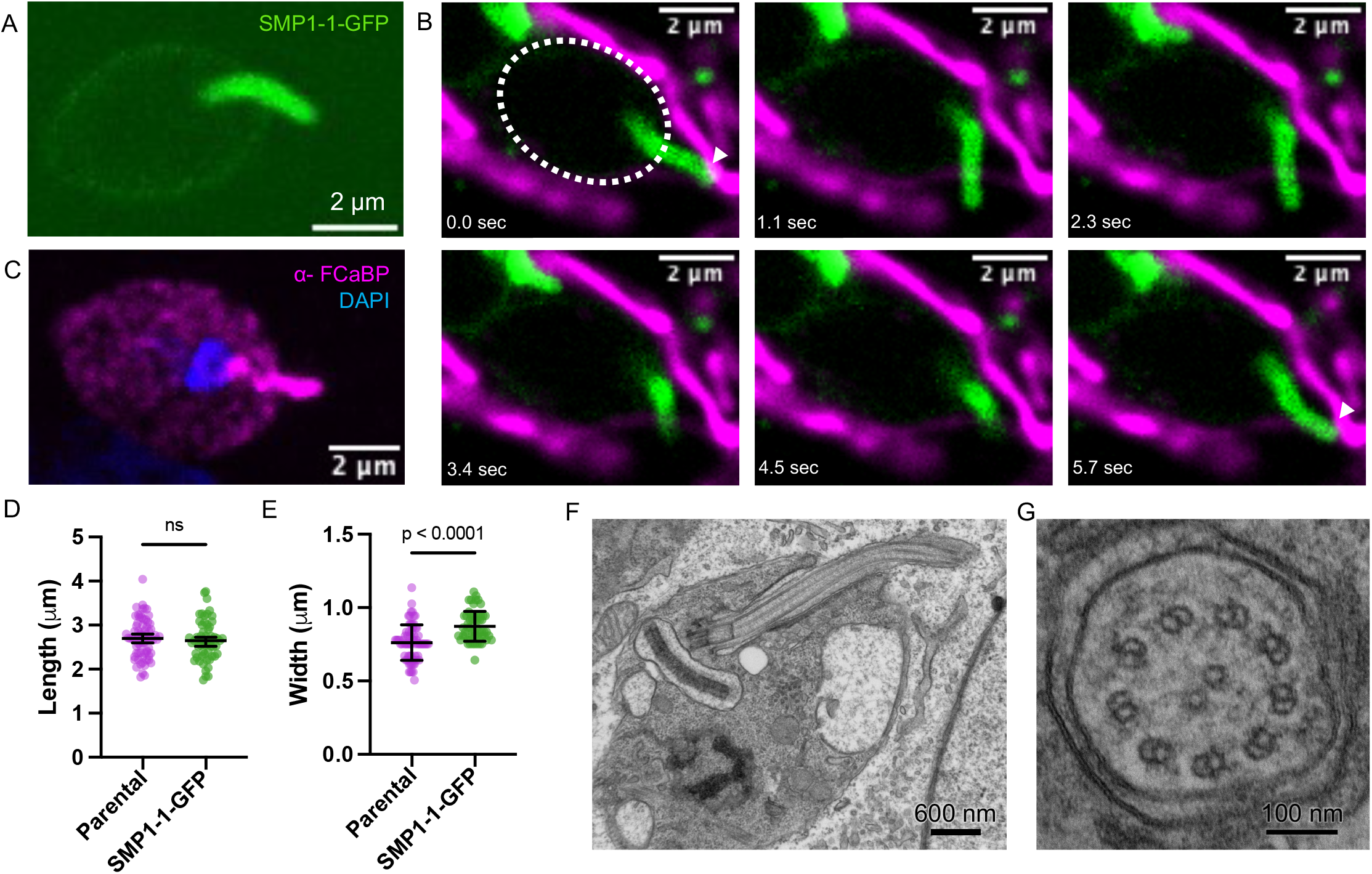
*T. cruzi* amastigotes have motile flagella. **A**: Live image showing a *T. cruzi* amastigote expressing SMP1-1-GFP. **B**: Time-lapse images of an SMP1-1-GFP*-*expressing *T. cruzi* amastigote (*green*) in the cytoplasm of a NHDF expressing mitochondrially-targeted mCherry (*pink*). The time each image was captured relative to the first frame in the series (arbitrarily set to 0.0 sec) appears at the top of each frame. White outline of amastigote body added on first frame, white arrows indicate points of contact. **C**: Image showing max projection of a z-stack from a *T. cruzi* amastigote inside of a NHDF cell, fixed with PFA and stained with α-FCaBP and DAPI. **D**: Flagellar length and **E**: width distribution of Tulahuén parental amastigotes fixed and stained at 48 hpi, as shown in C (n=64) in pink and SMP1-1-GFP expressing amastigotes imaged live in green. **F**: Electron microscopy of *T. cruzi* amastigote showing the flagellum longitudinally and **G**: in cross-section, showing a 9+2 axoneme.

In order to validate the use of SMP1-1-GFP expressing amastigotes as a tool for measuring the flagellar beat, we measured and compared the SMP1-1-GFP-expressing flagella (live imaging) with fixed parental flagella (immunofluorescence imaging), stained with an antibody to Flagellar Calcium Binding Protein (FCaBP, example image in **Fig 1C**) (Wingard et al., 2008). This analysis showed that an amastigote flagellum has an average length of 2.7 μm and an average width of 0.7-0.8 μm (**Fig 1D**, methodology **Fig S1**). The flagellar length and width measurements for the two populations were largely overlapping (**Fig 1D**) despite a slight decrease in flagellar width observed in fixed parental parasites (**Fig 1E**, ~0.1 μm), possibly due to shrinkage with fixation of the parental parasites. In comparison to the *Leishmania* amastigote, the flagellar length is ~1 μm longer, mostly accounted for by a longer external portion of the flagellum. Electron microscopy confirmed the length of the flagellar protrusion in the longitudinal section (**Fig 1F**). Examining the cross-section revealed a 9+2 axonemal structure (**Fig 1G**), the hallmark of motile cilia and flagella (Lindemann & Lesich, 2021), in agreement with previously published data (Gardener, 1974).

### Quantifying intracellular amastigote flagellar beats

To characterize *T. cruzi* amastigote flagellar movement, live imaging of SMP1-1-GFP-expressing intracellular amastigotes was performed at 48 hours post-infection of normal human dermal fibroblasts (NHDF). Time-lapse images captured for three independent infections were used to select twenty-four amastigotes from each replicate experiment for kymograph analysis. Dividing parasites (which feature two flagella) and those with morphological abnormalities were excluded from the analysis, although flagella of dividing amastigotes were clearly actively beating. Amira™ software was employed to create kymographs as 3D surface models of the fluorescence signal and track the course of the flagellar tip for a duration of 60 s. This allowed the recording of absolute coordinates for the amplitude of tip movement (example in **Fig 2A**). These data were then used to perform a peak analysis with OriginPro^®^ (example in **Fig 2B**). The distance between successive peaks was calculated and plotted to generate precise and normalized displacement measurements for each flagellum independent of the orientation or position of the parasite cell body. The frequency, temporal distribution, and displacement are plotted for each flagellar beat over the observation period of 1 minute for each parasite. Data obtained from five individual parasites are shown (**Fig 2C**), and plots for all 72 parasites are shown in **Fig S2. Figure 2D** is aggregated data for all parasites for each biological replicate. Although the flagellar beat frequency is highly variable on an individual parasite level, the average flagellar beat frequency is 0.69 ± 0.30 Hz. Such ‘quasiperiodic’ behavior can be expected in flagella shorter than 4 μm (Bottier et al., 2019). Note that all measurements are necessarily confined to the imaging plane and thus quantify the apparent microscopic behavior routinely captured in 2D. We observed clear indications of three-dimensional, rotational movement of the flagella but presume that the current detailed description is sufficient for an initial analysis of potential interactions in the host cytoplasm.

**Figure 2:**
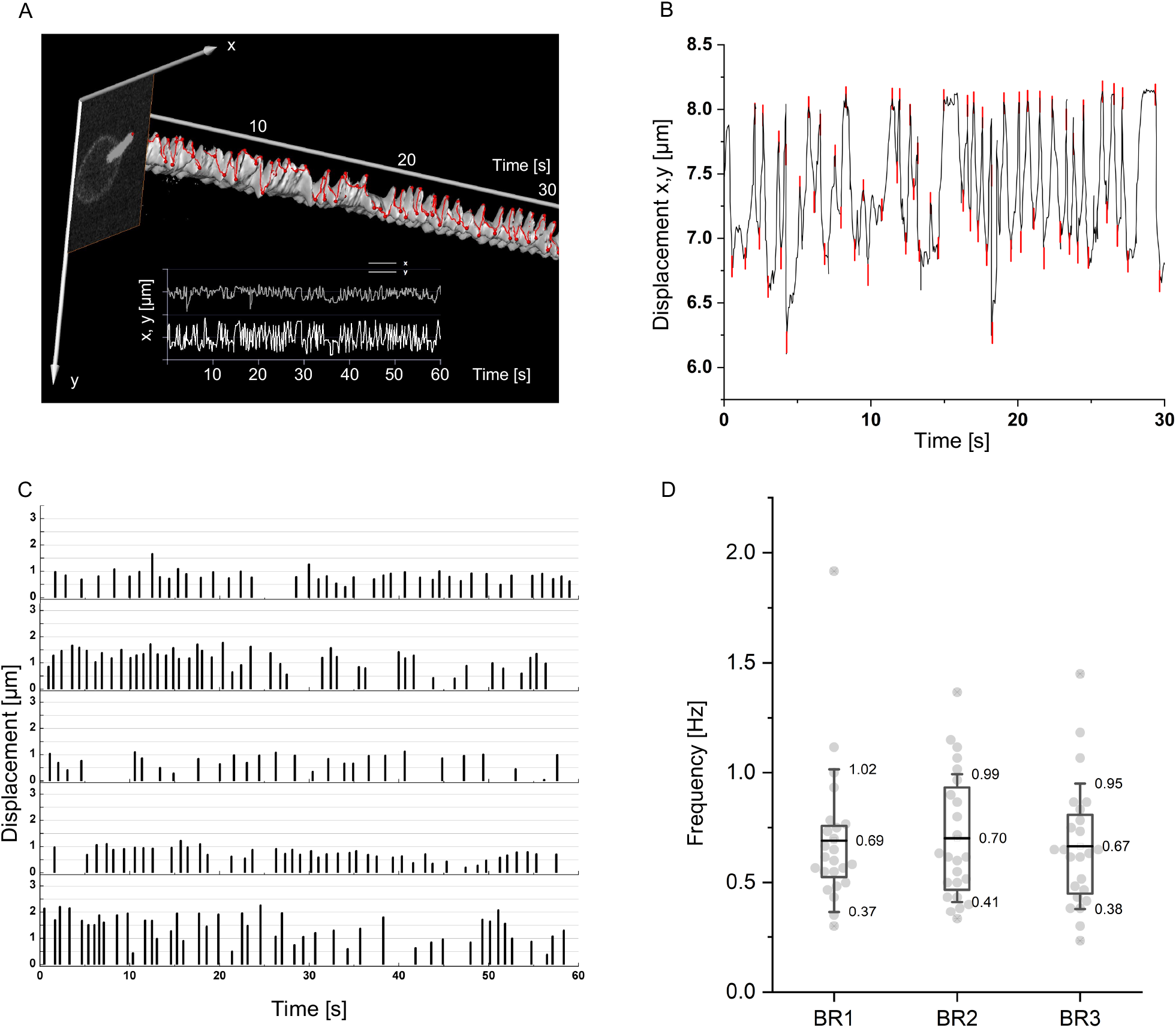
Quantifying intracellular amastigote flagellar beat. **A**: A 30-second example of the tip tracking trace generated using Amira™. **B**: A 30-second example of peak analysis in Origin^®^ with each peak shown in red. Baseline movement is subtracted in C. **C**: Data from 5 example parasites showing the number, amplitude (displacement), and temporal distribution of beats per minute. **D**: Aggregate data from all parasites from three biological replicates showing the flagellar beat frequency, each dot represents one parasite (n=24 per replicate).

### Flagellar movement is an intrinsic capacity of *T. cruzi* amastigotes and is impacted by calcium availability

As demonstrated above, flagellar movement is consistently observed for *T. cruzi* amastigotes that reside within mammalian host cells. To determine if the flagellar beat is an intrinsic property of amastigotes, SMP-1-1-GFP parasites were imaged following their release from mechanically disrupted host cells. We find that isolated *T. cruzi* amastigotes continue to beat their flagellum after separation from host cells and exhibit similar flagellar beat frequencies as host cell resident amastigotes, although slightly faster (~0.2 Hz, **Fig 3A**) potentially due to lowered viscosity outside of the cell. Furthermore, exposure of isolated amastigotes to GNF7686, an inhibitor of trypanosomatid cytochrome *b*, a component of the mitochondrial electron transport chain critical for ATP generation (Khare et al., 2015; Shah-Simpson et al., 2017), resulted in severe impairment of flagellar movement (**Fig. 3B**). The proportion of parasites with no measurable flagellar beat increased from 2% in untreated controls to 54% in the GNF7686-treated amastigotes (**Fig. 3B**). In line with the demonstration that GNF7686 lacks cytotoxicity toward *T. cruzi* amastigotes (Dumoulin et al., 2022), normal flagellar beating resumed in GNF7686-treated amastigotes upon washout of the compound (**Fig. 3B)**. Combined, these data support the conclusion that flagellar beating is an intrinsic property of *T. cruzi* amastigotes.

**Figure 3:**
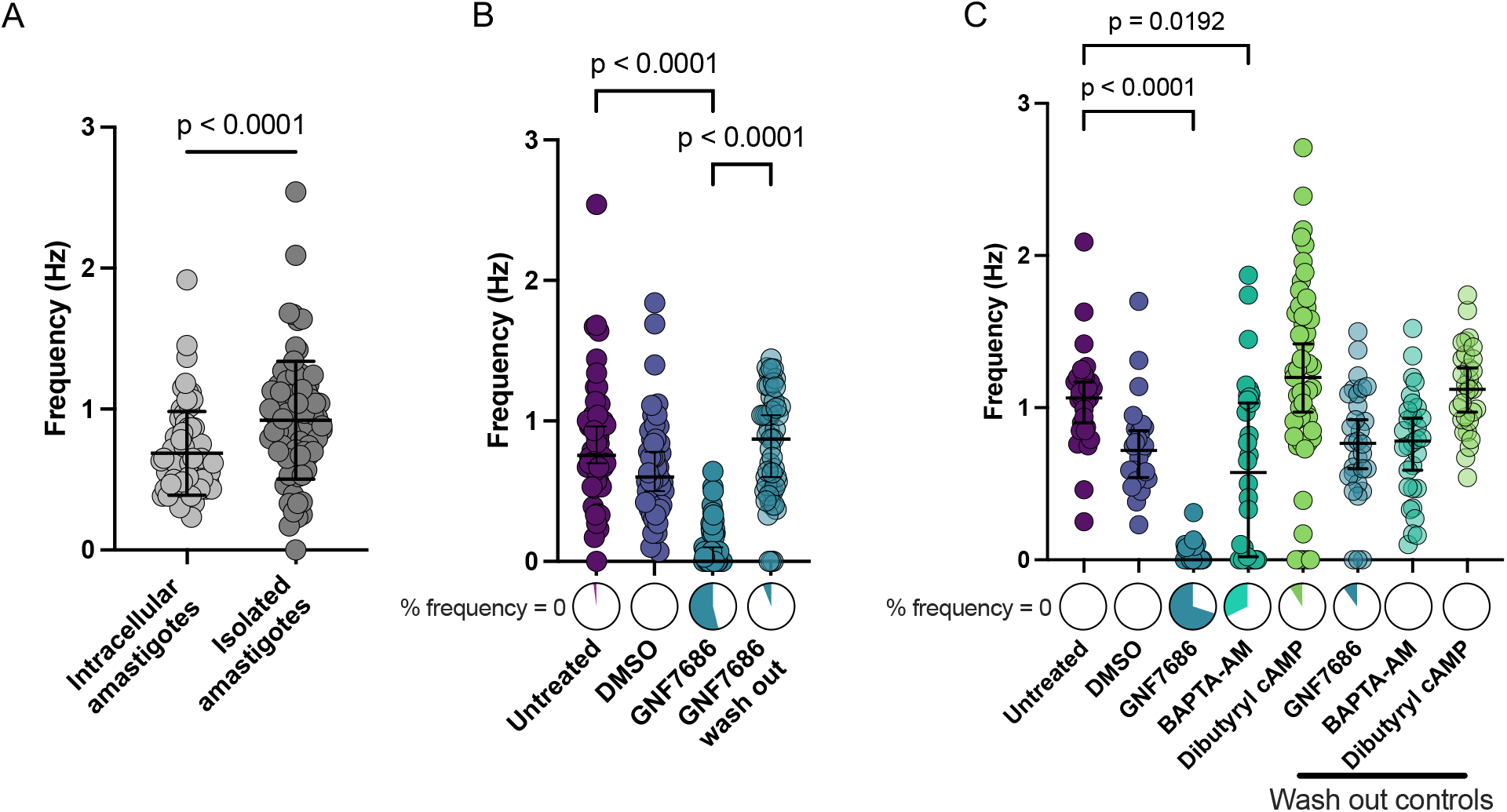
Intracellular movement is mirrored in extracellular environment and is dependent on amastigote metabolism and calcium availability. **A**: Frequency of flagellar beat for extracellular and intracellular amastigotes is shown (Mann-Whitney test). **B**: Parasite beat frequency of amastigotes treated with GNF7686, a parasite specific mitochondrial respiration inhibitor. Each treatment has a pre-treatment control, and the DMSO solvent control is shown. Results shown are the individual values for each parasite with mean and standard deviation lines. **C**: Flagellar beat frequency was assayed for amastigotes treated with GNF7686, BAPTA-AM, and dibutyryl cAMP, along with wash out controls. For B and C, pie charts above labels show percentage of parasite flagella that had a frequency of zero (did not move during observation period) in color. Statistical analysis of B and C was completed using a Kruskal Wallis test with Dunn’s multiple comparisons test.

Next, we used small molecules to perturb calcium and cyclic AMP homeostasis in *T. cruzi* amastigotes as these second messengers have been implicated in modulating eukaryotic flagellar beat and dynamics (Holwill & McGregor, 1976; Mukhopadhyay & Dey, 2016; Satarić et al., 2020). In related kinetoplastid species, proteins capable of binding calcium and cyclic AMP have been found in the flagellum and are implicated in modulating flagellar movement (Holwill & McGregor, 1976; Mukhopadhyay & Dey, 2016; Wu et al., 1992, 1994). We find that amastigotes treated with the membrane-permeant calcium chelator, BAPTA-AM, display decreased flagellar beat frequency, which is restored to near control levels upon washout of BAPTA-AM (**Fig 3C**). Exposure to dibutyryl cyclic AMP had no significant effect on average flagellar beat frequency (**Fig 3C**), although a trend toward higher variability in flagellar beat frequency was noted (10% non-motile flagella and 40% above 1.5 Hz) and recovered following washout of the compound.

Overall, this work has generated the first evidence of flagellar movement in the non-motile intracellular amastigote stage of a kinetoplastid parasite. To our knowledge, this is the first record of an intracellular eukaryotic flagellum beating within another eukaryotic cell. The biological role of amastigote flagellar beating is currently unknown, but it may serve as a dynamic sensor of metabolites in the host cytosol, potentially coming from the host mitochondria, given the previously observed interaction (Lentini et al., 2018). Alternatively, flagellar movement may facilitate endocytic uptake of cytosolic nutrients via the flagellar pocket (Langousis & Hill, 2014; Overath & Engstler, 2004; Soares & de Souza, 1991) or the cytostome-cytopharynx complex (Alcantara et al., 2021; Chasen et al., 2020). Finally, the intracellular beating flagellum could alter the physical properties of the host cell cytoplasm. It has been shown that the cytoplasm is a shear-thinning fluid (Rogers et al., 2008). Therefore, the flagellar beating will most likely increase the fluidity of the cytoplasm, which would significantly influence the diffusion dynamics and hence the molecular machinery of the host cell (Persson et al., 2020). Future studies will be aimed at defining the role of amastigote flagellar beating in the context of intracellular infection, including the dynamics of the unique amastigote flagellum-host mitochondrial interaction (Lentini et al., 2018).

## Materials and Methods Reagents

Compounds were purchased and diluted to stock concentrations: GNF7686 (Vitas-M Laboratory, Champaign, Illinois, USA), 5 mM stock in DMSO, Dibutyryl cAMP (Sigma, St. Louis, Missouri, USA), 200 mM stock in water, and BAPTA-AM (AAT Bioquest, Sunnyvale, CA, USA) 32.7 mM in DMSO.

### Mammalian and parasite cell culture

Mammalian cells were maintained in Dulbecco’s modified Eagle medium (DMEM; HyClone, Logan, Utah, USA) supplemented with 10% heat-inactivated FBS (Gibco, Waltham, Massachusetts, USA), 25 mM glucose, 2 mM L-glutamine, and 100 U/mL penicillin-streptomycin (DMEM-10) at 37°C and 5% CO_2_. *T. cruzi*-infected cultures infected with *Trypanosoma cruzi* were maintained in DMEM with 2% FBS (DMEM-2). Normal Human Neonatal Dermal Fibroblasts (NHDF; Lonza, Basel, Switzerland), mouse epithelial cells (MEF; ATCC, Manassas, Virginia, USA), LLC-MK2 (ATCC, Manassas, Virginia, USA) cells, and NHDF expressing a mitochondrially targeted mCherry (Lentini et al., 2018) were maintained as subconfluent monolayers and seeded the day prior to infection in all experiments.

*Trypanosoma cruzi* Tulahuén LacZ clone C4 was obtained from the American Type Culture Collection (ATCC, PRA-330; ATCC, Manassas, Virginia, USA). *T. cruzi* CL Brener, Y, and Brazil strains were obtained from Ricardo Gazzinelli (University of Massachusetts Medical School), Norma Andrews (University of Maryland), and Nisha Garg (University of Texas Medical Branch at Galveston), respectively. The axenic epimastigote stage of *T. cruzi* was propagated at 28°C in liver infusion tryptose (LIT) medium (4 g/L NaCl, 0.4 g/L KCl, 8 g/L Na2HPO4, 2 g/L dextrose, 3 g/L liver infusion broth, 5 g/L tryptose, with 25 mg/L hemin and 10% heat-inactivated FBS). After transfection, cloning, and selection of SMP1-1-GFP expressing parasites, they were converted to metacyclic trypomastigotes by incubating stationary phase epimastigotes in DMEM-2 for 5 days at 28°C. Cultures were washed in DMEM-2 and incubated with LLC-MK2 cells at 37°C in a 5% CO_2_ incubator to allow host cell invasion. Mammalian stage trypomastigotes that emerged from infected LLC-MK2 cells were harvested from the culture supernatant and used to infect fresh cells. This cycle was continued on a weekly basis to maintain the mammalian-infective cycle. Trypomastigotes were prepared by collecting the supernatant from infected cultures and centrifuging for 10 minutes at 2060 x g, then either the trypomastigotes were allowed to swim up from the pellet and collected again, or they were immediately washed in DMEM-2 and quantified for subsequent infections.

### *T. cruzi* strain generation

*T. cruzi* epimastigotes were transfected with a modified pTREX plasmid (DaRocha et al., 2004; Dumoulin et al., 2022) containing SMP1-1 and eGFP (Cormack et al., 1996). The full-length SMP1-1 sequence (TcCLB.506563.200) was PCR amplified from genomic DNA isolated from *T. cruzi* Tulahuén strain parasites using the following primers: 5’-ATCGTAGAATTCATGGGCTGCGGTGCTTCTTCGAAACCCTCCAC-3’ 5’-ATCGTAGCGGCCGCATTTTCTTTTCTTCTTCTTCCTTCGGGCGTTTGTTTTTTTCAGTGGGGACGGC-3’ Prior to transfection, epimastigotes were pelleted at 2060 x g for 10 minutes, then resuspended in 100 μL of Tb BSF buffer (Schumann Burkard et al., 2011). 4×10^7^ epimastigotes were loaded into a sterile 2 mm gap cuvette and transfected using an Amaxa Nucleofector II (Lonza, Basel, Switzerland; U-33 program). Parasites were immediately transferred to LIT medium for 24 hours before adding 10 μg/mL puromycin (Invivogen, San Diego, California, USA). After drug selection, SMP1-1-GFP expressing Tulahuén epimastigotes were cloned by limiting dilution in 96-well plates, and a single clone was selected for experimental use based on >90% SMP1-1-GFP expression. For the other parasite strains (Brazil, Y, and CL Brener), uncloned populations of SMP1-1-GFP expressing parasites were used for imaging.

### Electron microscopy

At 48 hpi, infected NHDF grown on Aclar (Ted Pella Inc., Redding, California, USA) filmed plastic coverslips were fixed with 1.25% formaldehyde, 2.5% glutaraldehyde, and 0.03% picric acid in 0.1 M sodium cacodylate buffer, pH 7.4 for 1 hour, and then washed 3 times in 0.1 M sodium cacodylate buffer (pH 7.4) prior to the post-fixation processing step of 1% osmium tetroxide/1.5% potassium ferrocyanide in distilled water 30 min on ice. Following three washes in distilled water, coverslips were incubated overnight with 1% aqueous uranyl acetate at 4 °C in the dark. Samples were rinsed in water and dehydrated in a graded ethanol series using the progressive lowering of temperature method. After a final dip in fresh 100% ethanol and then 100% propylene oxide, they were infiltrated with solutions 2:1, 1:2 of propylene oxide:epon araldite 30 min each, then 100% Epon araldite for 1 hour, then mounted for polymerization at 65 °C for 48 hr. Ultrathin sections (about 60 nm) were cut on a Reichert Ultracut-S microtome (Leica, Wetzlar, Germany), picked up on to copper grids stained with lead citrate, and examined in a TecnaiG2 Spirit BioTWIN (FEI Company, Hillsboro, Oregon, USA). Images were recorded with an AMT 2k CCD camera (Advanced Microscopy Techniques, Woburn, Massachusetts, USA).

### Indirect immunofluorescence microscopy and flagellar morphology quantification

NHDF cells were seeded onto round glass coverslips (12 mm, #1.5; Electron Microscopy Sciences, Hatfield, Pennsylvania, USA) in 24-well plates at a density of 20,000 cells/well in DMEM-10 at 37°C, 5% CO_2_. Cells were infected the following day with WT Tulahuén trypomastigotes at a multiplicity of infection (MOI) of 3 suspended in DMEM-2 and washed twice with Phosphate Buffed Saline (PBS; Corning, Corning, New York, USA) 24 hours after infection and incubated further in DMEM-2 at 37°C, 5% CO_2_. At 48 hours post-infection (hpi), the media was replaced with a 1% paraformaldehyde solution in PBS for a 10-minute incubation at 4°C. All the following steps were carried out at room temperature, and each was preceded by three washes of the wells with PBS. Cells and parasites were permeabilized with 0.1% Triton-X 100 (v/v) (JT Baker, Phillipsburg, New Jersey, USA) for 10 minutes and a blocking solution of 3% (w/v) Bovine Serum Albumin (Sigma-Aldrich, St. Louis, Missouri, USA) in PBS for 1 hour. The primary antibody solution containing 1:1,500 rabbit α-FCaBP (Maric et al., 2015) in 1% BSA in PBS was added for 1 hour, followed by a 1:1000 α-Rabbit Alexa Flour 647 solution in 1% BSA in PBS for 1 hour. DAPI (0.2 μg/mL; Thermo Fisher Scientific, Waltham, Massachusetts, USA) in PBS was added for 5 minutes, and following washes, coverslips were placed onto slides with Prolong^®^ Diamond mounting medium (Thermo Fisher Scientific, Waltham, Massachusetts, USA). After setting for 24 hours, the cells were imaged with a 100x objective using a Yokogawa CSU-X1 spinning disk confocal system paired with a Nikon Ti-E inverted microscope and an iXon Ultra 888 EMCCD camera. Image processing, analysis, and display were completed using FIJI (Schindelin et al., 2012). Complete image analysis methods and an example are presented in **Fig S1**.

### Amastigote isolation

NHDF cells were seeded at 3 × 10^7^ cells per T-150 flask and infected with SMP1-1-GFP expressing *T. cruzi* trypomastigotes (MOI of 20) in DMEM-2 at 37°C, 5% CO_2_ for 18 hr. The remaining extracellular parasites were removed, and infected cells were further incubated at 37°C, 5% CO_2_. At 48 hpi, infected monolayers were washed twice with PBS, and cells were dissociated using Accumax (Innovative Cell Technologies, San Diego, California, USA). Cell suspensions were washed 2 times in PBS, loaded into gentleMACS M tubes (Miltenyi Biotec, Cologne, Germany) with a total volume of 2.5 mL, and cells were lysed using the Protein_01.01 M tube protocol on the gentleMACS Dissociator (Miltenyi Biotec, Cologne, Germany). Amastigotes were pelleted at 2060 x g for 10 minutes, then resuspended in imaging medium at a final concentration of 1.45 ×10^7^ amastigotes per milliliter.

### Live cell confocal microscopy

*Intracellular amastigotes*: 30,000 NHDF were seeded onto 35 mm glass bottom dishes (Matsunami Glass, Bellingham, Washington, USA) in DMEM-10 and allowed to attach for 24 hr at 37°C, 5% CO_2_. Cells were incubated with untagged or SMP1-1-GFP expressing *T. cruzi* trypomastigotes (MOI between 3 and 30) for 24 hr in DMEM-2 at 37°C, 5% CO_2_ to allow infection. The remaining extracellular parasites were removed by rinsing monolayers followed by further incubation in DMEM-2. At 48 hpi medium was aspirated and replaced with imaging medium: Fluorobrite™ DMEM (HyClone, Logan, Utah, USA) supplemented with 1.5% FBS, 2 mM glutamine, and ProLong™ Live Antifade Reagent (1:75; Invitrogen, Waltham, Massachusetts, USA). Dishes were imaged using a Yokogawa CSU-X1 spinning disk confocal system paired with a Nikon Ti-E inverted microscope equipped with a 37°C CO_2_ injectable environmental chamber and an iXon Ultra 888 EMCCD camera. All images were acquired with the 100x objective with a total imaging time between 30 seconds and 1 minute. Temporal resolution for intracellular amastigotes is 30 ms for the Tulahuén strain and 200 ms for the other strains in the supplemental videos. Dual channel imaging (**Supplemental Video 1**) has a temporal resolution of 1.08 s.

*Isolated amastigotes:* 2.2 × 10^6^ freshly isolated intracellular *T. cruzi* amastigotes in 150 μL of pre-warmed imaging medium were placed directly onto the glass coverslip of a 35 mm glass bottom dish and allowed to settle at 37°C in a 5% CO_2_ incubator for 10 minutes. 75 μL of media was removed prior to placing the dish on the microscope for imaging. For small molecule treatments, 150 μL of isolated amastigotes were exposed to the compound at the indicated final concentrations for 10 min, then either directly transferred to the imaging well and allowed to settle for 10 min as above or for compound washout experiments, treated parasites were diluted 100-fold in PBS and centrifuged at 2060 x g for 10 minutes before resuspending in 150 μl imaging medium and proceeding as above. Temporal resolution for intracellular amastigotes is 80 ms.

### Quantification of flagellar beat

*Intracellular amastigotes*: Kymographs of fluorescence microscopy time-lapse series were analyzed using Amira (2019.1, Thermo Fischer Scientific). Recordings of fluorescence-labeled flagella (SMP1-1-GFP) with a duration of 60 seconds (s) and a temporal resolution of 30 ms were used to create surface models using the timescale as the z-axis, thus creating 3d-kymographs of the fluorescence signal (e.g., **Fig. 2A**). Each surface model was rotated to a position in which the movement of the flagellum tip was clearly visible as an oscillating pattern on the surface model. The flagellar tip movement was then traced manually along the surface, creating a surface geodesic path (red track, **Fig. 2A**), yielding the coordinates of the tip in the plane of the original images for 60s time periods. The measurements of the x and y positions quantified the oscillations of the flagellum during the 60s recording.

To generate a standardized and normalized dataset for the evaluation of beat frequency and amplitude, the x, y positions were exported and analyzed using the peak analysis tool of OriginPro (2021, OriginLab). The x and y values were converted into vector coordinates for peak identification in the positive and negative direction (e.g., **Fig. 2B**). The Euclidian distance between two successive peaks of opposite direction was then calculated, yielding the distance covered by the flagellum tip during each beat. This measurement of tip movement is independent of the cell orientation in the 2d image plane and thus normalizes the apparent amplitude of the flagellar beat (**Fig. 2C**). The tip movement between two peaks was significantly larger than movements of the tip caused by the diffusional displacement of the entire cell in the same time period, which could therefore simply be thresholded in the peak detection step. The average frequency [Hz] was calculated by dividing the number of measured beats by 60 [s].

#### Isolated amastigotes

The frequency of the extracellular amastigote flagellar beat was determined by counting the number of beats during visual inspection of the 60s time-lapse series. A beat was defined by an obvious bending of the axoneme, causing the flagellar tip to move a distance significantly further than any displacement of the cell body in the same time period, equivalent to the criteria used in peak analysis (see above). Statistical analysis was completed, as noted in figure legends using GraphPad Prism version 9.4.1 for Mac (GraphPad Software, San Diego, California, USA).

## Acknowledgments

We thank the Sabri Ülker Center’s Advanced Imaging Lab at the Harvard T.H. Chan School of Public Health for microscopy training and access. We thank Ricardo Gazzinelli (University of Massachusetts Medical School), Norma Andrews (University of Maryland), and Nisha Garg (University of Texas Medical Branch at Galveston) for parasite strains and David Engman (Cedars-Sinai Medical Center) for the gift of the FCaBP primary antibody. We also thank Lucas Pagura for his thoughtful discussion and Gaelle Lentini for her foundational work supporting this project. This work was supported by NIH grant R21AI135520 awarded to B.A.B. and the 5T32AI049928 T32 training grant. M.E. was funded by the Deutsche Forschungsgemeinschaft (DFG grants EN305; SPP1726; SPP2332; GRK2157)), the German Federal Ministry of Education and Research (BMBF, NUM Organostrat), the European Community (EU, Phymot) and the German-Israeli Foundation for Scientific Research and Development (GIF, I-473-416.12/2018).

