## Supplemental Figures for "The intracellular amastigote of *Trypanosoma cruzi* maintains an actively beating flagellum"

Supplemental Figure 1

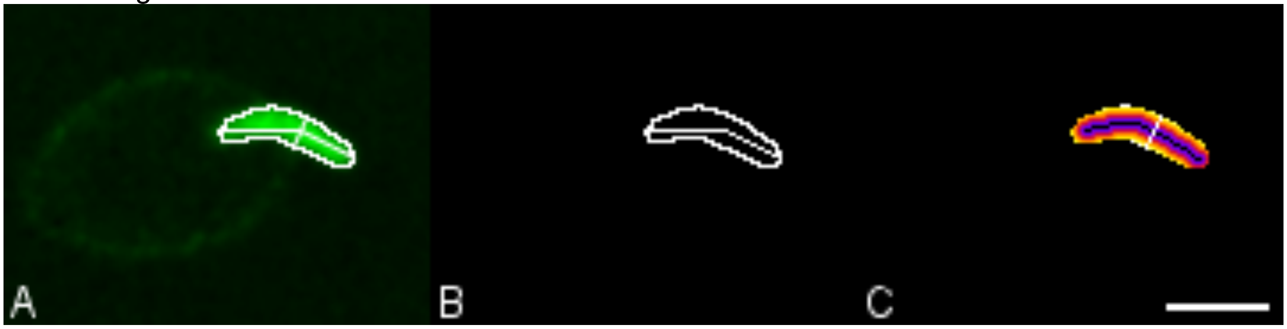


**Supplemental Figure 1: Method for automated length and width measurement of fluorescent flagella using Fiji including the MorphoLibJ plugin. A:** Selected image of focused flagellum showing length and width measurements (Fig. 1C: l = 3,03 µm, w = 0,97 µm). **B:** Flagellum labelled by “Analyze Particles” after image A was binarized using “Auto threshold/Method Yen”. The dashed line shows the “Geodesic Diameter” as determined by the MorphoLibJ plugin and used for length measurement. **C:** Geodesic distance map (MorphoLibJ) of the binarized image in B after using the plugin “Skeletonize” in order to determine the flagellum´s midline (black line). The highest value of the distance to the midline (colour coded bright yellow) was used to calculate the width of the flagellum at the widest position (white line).

Supplemental Figure 2

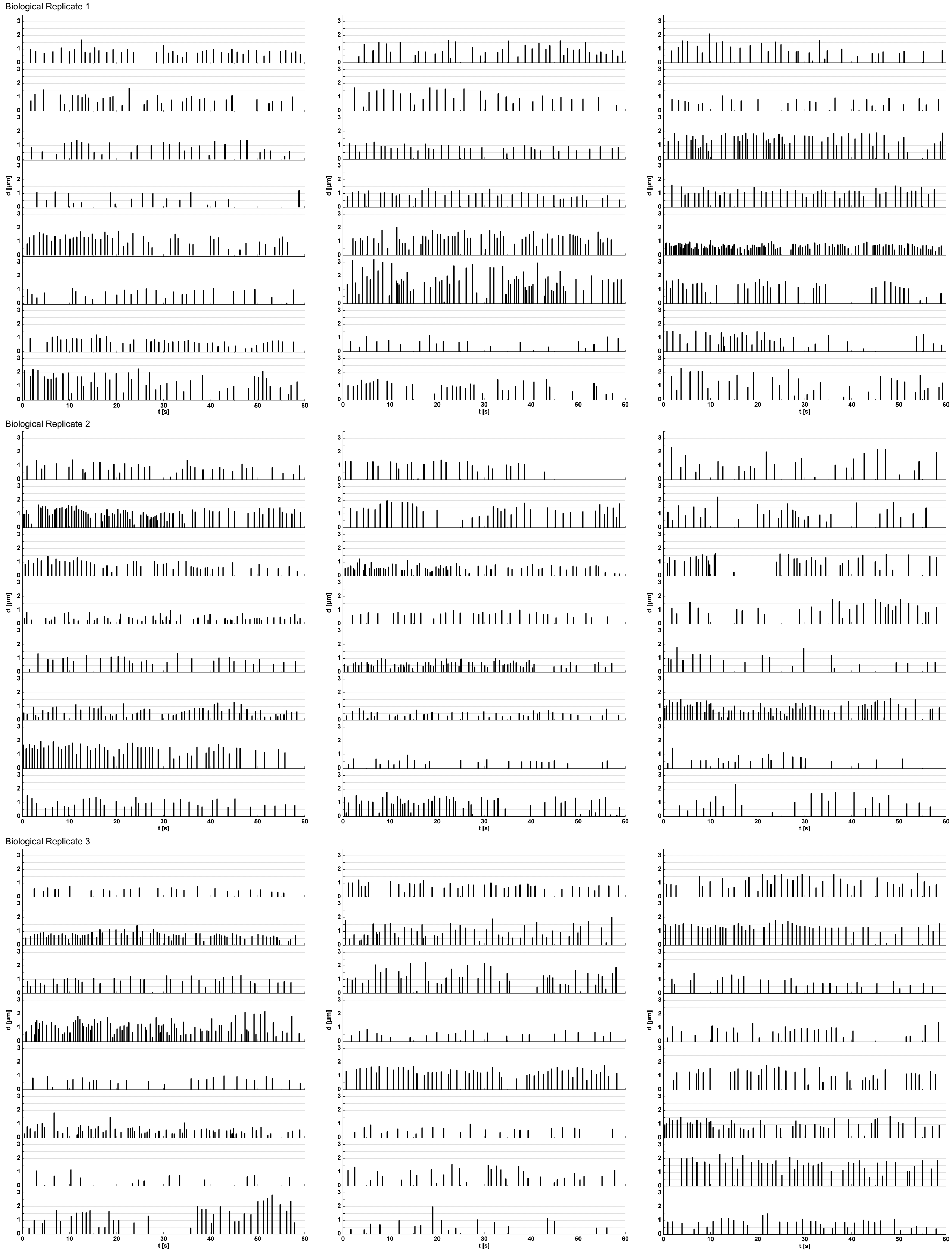


**Supplemental Figure 2: Individual intracellular amastigote flagellar beat profiles**. Flagellar beat profiles showing the number, amplitude, and temporal distribution of beats per minute for all parasites in Fig 2D.
