## Supplemental Video Legends for "The intracellular amastigote of *Trypanosoma cruzi* maintains an actively beating flagellum"

**Supplemental video 1**: Live cell imaging of a *T. cruzi* Tulahuén strain amastigote expressing SMP1-1-GFP inside of a normal human neonatal dermal fibroblast.

**Supplemental video 2**: Live cell bright-field imaging of *T. cruzi* Tulahuén strain amastigotes inside of a normal human neonatal dermal fibroblast.

**Supplemental video 3**: Live cell imaging of a *T. cruzi* CL Brener strain amastigote expressing SMP1-1-GFP inside of a normal human neonatal dermal fibroblast.

**Supplemental video 4**: Live cell imaging of a *T. cruzi* Y strain amastigote expressing SMP1-1-GFP inside of a normal human neonatal dermal fibroblast.

**Supplemental video 5**: Live cell imaging of a *T. cruzi* Brazil strain amastigote expressing SMP1-1-GFP inside of a normal human neonatal dermal fibroblast.

**Supplemental video 6**: Live cell imaging of a *T. cruzi* Tulahuén strain amastigote expressing SMP1-1-GFP inside of a mouse epithelial cell.
